# Cell-extrinsic effects in T cell acute lymphoblastic pre-leukemia stem cells mediated by EphA3

**DOI:** 10.1101/2020.09.14.297127

**Authors:** Adriana C. Pliego Zamora, Hansini Ranasinghe, Jessica E. Lisle, Stephen Huang, Racheal Wadlow, Andrew M. Scott, Andrew W. Boyd, Christopher I. Slape

**Author notes:** Correspondence to: Christopher I. Slape, The University of Queensland Diamantina Institute, Level 5, TRI Building, 37 Kent St, Woolloongabba QLD 4102 Australia.

## Abstract

Our recent study of a novel model of T-ALL pre-leukemic stem cells, the NUP98-HOXD13 (NHD13) mouse, showed that the abnormal self-renewal of these stem cells was dependent on Lyl1 yet, when Lyl1 was deleted, the T-ALL still developed. In the present study, we observe that the thymocytes in these mice also overexpress EphA3, and we characterise the thymocytes in NHD13-EphA3^−/−^ mice. NHD13-EphA3^−/−^ thymocytes retain their abnormal self-renewal activity demonstrated by their capacity to engraft following primary and secondary transplants. Strikingly, NHD13-EphA3^−/−^ thymocytes fail to engraft upon the third serial transplant, whereas the NHD13 thymocytes engraft indefinitely. Seeking to explain this, we find that NHD13 DN2 thymocytes are capable of halting the normal differentiation process of incoming WT progenitor cells, and remarkably, this capacity is severely impaired in the absence of EphA3. Therefore EphA3 is not critical for engraftment, but is essential for enabling the halt in differentiation of neighbouring WT cells, which in turn allows the incumbent progenitors to remain longer in the thymus due to an absence of normal cell competition, a property that in itself has been demonstrated to be oncogenic. We suggest that pre-leukemic self-renewal in this model is a complex interplay of cell intrinsic and extrinsic factors, and that multiple redundant pathways to leukemogenesis are active in this model.

## Introduction

T-ALL is a clonal malignancy caused by the accumulation of genomic lesions that disrupt the development of T cells. According to the ectopic expression of basic helix-loop-helix transcription factors, LIM domain proteins, homeobox transcription factors and HOXA gene clusters, T-ALL cases can be divided into subfamilies including: TAL/LMO, immature/early T-cell precursor (ETP-like), TLX1/TLX3 or HOXA cluster, respectively [1–8]. Current therapy for T-ALL is non-targeted chemotherapy, which achieves disease-free status in the majority of cases, especially in paediatric patients [9, 10]. However, relapse can still occur and is often refractory to treatment and fatal. Relapse is caused by the presence of leukemic stem cells (LSC) that are resistant to standard treatment [11, 12]. The hallmark of LSCs is that these cells can self-renew, proliferate and differentiate [1, 13–18].

The Nup98-HoxD13 (NHD13) transgenic mouse is a model of HoxA-driven T-ALL, with pre-leukemic thymocytes that exhibit LSC-like features before the overt development of T-ALL. Specifically, NHD13 thymocytes self-renew (ie engraft upon transplantation), accumulate in the DN2 differentiation stage and, in one third of cases, lead to the development of T-ALL [19–21]. NHD13 thymocytes require the Lyl1 gene for self-renewal, but although NHD13 Lyl1 knockout (KO) thymocytes cannot engraft, these mice still succumb to T-ALL [21]. Therefore, it appears that the ability of NHD13 thymocytes to engraft is not required for the induction of T-ALL, and other mechanisms must be considered.

Another potential mechanism of formation of T-ALL is an absence of cell competition in the thymus. In health, T lymphocytes are continually seeded in the thymus, migrating from the bone marrow, throughout life. These progenitors differentiate through a series of well-defined stages in the thymus before entering the circulation as mature T-cells [22, 23]. Incoming progenitors compete out incumbent progenitors, ensuring regular turnover. Recent experiments have shown that in the absence of incoming progenitors and therefore cell competition, incumbent thymocytes can aberrantly self-renew. These long-lived thymocytes, however, leave the organism predisposed to T cell-acute lymphoblastic leukaemia (T-ALL) [23–28]. Normal cell competition, therefore, is a tumour suppressor in the thymus [23, 25]. We considered a reduction in cell competition as a possible alternate mechanism by which T-ALL could be induced in NHD13 mice.

The Eph family of Receptor tyrosine kinases and their ligands, ephrins, can modulate cell adhesive properties as well as coordinate cell movement thus playing critical roles in development, tissue homeostasis and cancer [29–38]. It has been hypothesized that this family of RTKs could also regulate CSCs functions [39]. Importantly, EphA2 is important for tumour suppressive cell competition in epithelial sheets [40]. Of particular interest, EphA3 is overexpressed in many different types of leukemias including T-ALL [41–45]. There is evidence implicating EphA3 in regulating stem-like features in LSC [46] and solid CSC [47]. EphA3 is a promising therapeutic target [48], especially in glioblastoma [49, 50]. Moreover, EphA3 is a viable anti-leukemic target, with the success of an activating monoclonal antibody (IIIA4), targeted payload delivery, or RNAi-mediated EphA3 knockdown, in models of multiple myeloma [51] and pre-B-ALL [52] as well as in human acute myeloid leukaemia (AML) patients [44].

In this study, we used NHD13 thymocytes as a model of pre-LSC to investigate the possible role of EphA3 in the development of T-ALL. We found that NHD13 thymocytes aberrantly express EphA3 and that this gene is not essential for short-term engraftment following transplantation. However, loss of EphA3 restores normal cell competition in the thymus, leading to loss of transplantation by the third passage. Unlike the loss of Lyl1, loss of EphA3 reduces the incidence of T-ALL. We suggest that EphA3 blocks cell competition in this model, and that this mechanism can explain the continued incidence of T-ALL in the absence of Lyl1.

## Methods

### Mouse strains

Transgenic NHD13 mouse [20] and EphA3KO mouse [53] have been previously described. The double transgenic NHD13-EphA3^−/−^ mouse was generated by crossing NHD13 and EphA3KO mice. All strains were back crossed with a C57BL/6 background. CD45.1 WT mice were purchased from ARC, Perth Australia. Mice were euthanized at 6 or 12 week old accordingly, using CO_2_ then thymuses were dissected. Animal experiments were approved by the Animal Ethics Committee of the University of Queensland.

### Tissue processing

Single-cell thymocytes suspensions were obtained by gently pressing the thymus onto a 40 µm cell strainer (BD Falcon) with a syringe plunger and rinsing with PBS supplemented with 2% FBS (2% FBS-PBS). Cell counts and cell viability were determined using Trypan blue. Thymocytes samples were used for flow cytometry analysis, transplantation experiments or stored in TRIzol (Invitrogen) at −80°C.

### Flow cytometry: Thymocyte DN populations, chimerism and cell cycle

For each mouse 1×10^6^ thymocytes were blocked with anti CD16/32 (FcR receptor) and stained with Zombie Aqua, lineage (B220, Ter119, Gr-1, Mac-1), CD4, CD8, CD44, CD25, C-kit and EphA3. Double negative (DN) populations were determined as lineage^−^, CD4^−^, CD8^−^, CD44^−/+^, CD25^−/+^ and C-kit ^−/+^. The expression of EphA3 was evaluated in each DN 1-4 populations. In addition to the DN panel, CD45.1 and CD45.2 were included for chimerism studies. For cell cycle studies, cells were stained as previously described for extracellular markers, then fixed and permeabilized with Fix & Perm Cell Permeabilization reagents (Thermo Fischer), stained first with Ki67 and then with 20µM Hoechst 33342 (Sigma Aldrich/Merck) in a solution with 0.05% Saponin (Sigma Aldrich/Merck) and 1µg/ml RNaseA (Sigma Aldrich/Merck) [54]. Cells were analysed by flow cytometry using a BD LSR FortessaX-20. Data was analysed using FlowJo 10.

### Serial transplantations

6 week old donors (CD45.2) of each indicated genotype were euthanised using CO_2_ and thymuses were dissected. Single thymocyte suspension was obtained as previously described. One sixth of the thymus in 200 µl of PBS was intravenously injected into female mice (CD45.1) sub-lethally irradiated with 6.5 Gy from a ^60^C γ. Four weeks post-transplantation, recipients were euthanised and a serial transplantation was performed as described above for a total of 4 subsequent times. Samples of thymocytes were stained and analysed for chimerism and cell cycle studies, as previously described.

### qRT-PCR

Total RNA was isolated from thymocytes using TRIzol (Invitrogen) reagent and GenElute mammalian total RNA miniprep kit (Sigma Aldrich). Day 17 embrionic RNA was used to validate primers (Clontech/Takara). cDNA was synthesised with Superscript III (Invitrogen). Primers for HoxA5, HoxA7, HoxA9, HoxA10, Lmo2, Hhex and Klf2 were designed in house and purchased from IDT. For all qRT-PCR reactions, PowerUp™ SYBR™ Green Master Mix (Applied Biosystems) was used, except for Lyl1 Taqman Fast Advanced Master Mix (Applied Biosystems) was used. Relative expression of target transcripts was analysed on ViiA 7 Real-Time PCR System (Thermo Fischer Scientific). Expression was normalised to two reference genes, HPRT1 and B2M.

### Survival cohort

Mice groups were designed to enable detection of a relative hazard of 0.5 with type I and II error rates of 0.3. The presented data excludes myeloid leukemias and myelodysplatic cases. Mice that reached ethical end-points were euthanised using CO_2_ and thymomas/thymuses were harvested. T-ALL cell profiles were used to corroborate diagnosis. Single thymocytes samples were obtained, stained and analysed by flow cytometry as previously described.

### Statistical Analyses

GraphPad Prism 7.3. was used for analysis. One-way ANOVA with Tukey’s Post-hoc analysis. Survival curve analysis was performed by the Mantel-Cox test. *p<0.03, **p<0.002, ***p<0.0002 and ****p<0.0001. Bars represent the SEM.

## Results

### EphA3 is abnormally expressed in NHD13 thymocytes

We characterised the cell profiles of WT and NHD13 thymuses by flow cytometry (Fig.1A). Consistent with previous studies [21], the NHD13 thymuses showed an abnormal accumulation of thymocytes in the DN2A (CD44^hi^, CD25^+^) stage (Fig.1B). We also observed an abnormally high level of expression of the c-kit protein in the CD25^+^ fraction (Fig. 1B) as previously described [21]. We examined the expression of EphA3 by flow cytometry and found that EphA3 is specifically expressed in the DN2A (CD4^−^, CD8^−^, CD44^+^, CD25^−^) self-renewing population of NHD13-EphA3^+/+^ thymocytes (Fig.1C).

**Figure 1.**
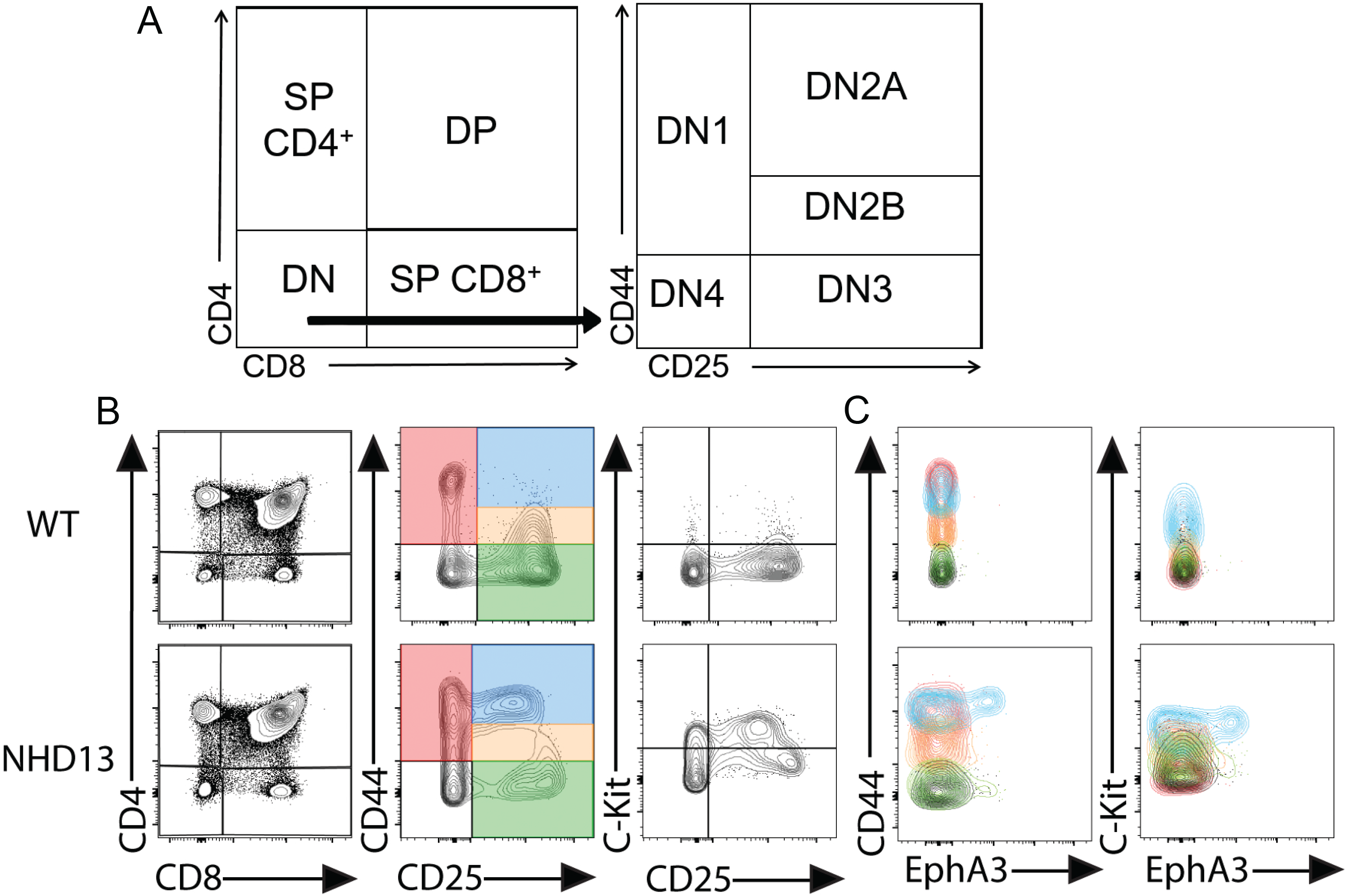
EphA3 is expressed in DN2A NHD13 thymocytes. **A)** Gating strategy to examine thymocytes by flow cytometry. **B)** Representative cell profiles using CD4/CD8, CD44/CD25, C-Kit/CD25. **C)** Detection of EphA3 in DN1-4. The colour of the contour plots matches the colour of the gated DN1-4 populations from panel B. NHD13 thymocytes present an abnormal differentiation block in the DN2A (CD44^hi^, CD25^+^, C-Kit^+^) differentiation stage.

### Characterisation of NHD13 EphA3 knockout mouse

To assess the role of EphA3 in the NHD13 thymus, we crossed the NHD13 transgene onto an EphA3 deficient background to generate EphA3-deficient NHD13 mice (NHD13-EphA3^−/−^). We confirmed the absence of EphA3 on the thymocytes by flow cytometry (Fig. 2A). Analysis of the thymuses of each genotype (WT, EphA3^−/−^, NHD13-EphA3^+/+^ and NHD13-EphA3^−/−^) at 6 weeks of age showed the CD4/CD8 cell profiles (Fig. 2B) and proportions of DN (CD4^−^/CD8^−^), DP (CD4^+^/CD8^+^), SP4 (CD4^+^/CD8^−^) and SP8 (CD4^−^/CD8^+^) populations were the same (Supp. Fig. 1). However, there were significant differences in the cellularity of the thymus and the proportions of the DN (CD44/CD25) sub-populations. Notably, the low cellularity was normalised (Fig. 2D) and the accumulation of DN2A cells was decreased (Fig. 2B and 2D) in the NHD13-EphA3^−/−^ thymuses. The overexpression of C-kit remained abnormally high in both NHD13-EphA3^+/+^ and NHD13-EphA3^−/−^ DN2A thymocytes. The proportions of the DN1 and DN2B populations in the NHD13-EphA3^−/−^ thymus are similar to those in the WT and EphA3^−/−^ control thymuses, correcting the abnormalities seen in the NHD13-EphA3^+/+^ thymus. The proportion of DN2A and DN3 cells was also partially normalised (Fig. 2D). Yet, the distribution of the DN profile of the NHD13-EphA3^−/−^ was unlike those of any of the other genotypes (Fig. 2B), suggesting that despite their similarities, a biological difference between the NHD13-EphA3^−/−^ and the WT and EphA3^−/−^ controls remains. To gain further insight, we characterised a cohort of 12 week old mice. Interestingly, at 12 weeks of age the loss of EphA3 did not rescue the phenotype of NHD13 thymuses, as shown by the similarity in CD4/CD8 cell profiles (Fig. 2C) and significantly increased DN, SP4 and SP8 populations as well as decreased DP populations (Supp. Fig. 1) in both the NHD13-EphA3^+/+^ and NHD13-EphA3^−/−^ thymuses. Moreover, the low cellularity (Fig. 2E), accumulation of DN2A thymocytes (Fig. 2C and E), increased DN1 and decreased DN3 proportions (Fig. 2E) are the same in both NHD13-EphA3^+/+^ and NHD13-EphA3^−/−^ thymuses. Therefore, phenotypes that were rescued or partially rescued at 6 weeks of age were once again abnormal at 12 weeks of age. In the thymus of WT mice, the loss of EphA3 had no effect at either time point, exhibited by no significant changes in the thymus cellularity (Fig. 2D and E), equal proportions of DN, DP, SP4, SP8 (Supp. Fig. 1) and DN1-4 (Fig. 2D and E) populations, as well as equal cell profiles (Fig. 2B and C) in both WT and EphA3^−/−^. Overall, these results suggest that the deletion of EphA3 transiently rescues the NHD13 thymus phenotype, displaying a WT-like phenotype at early time points that reverts to an NHD13-like phenotypes at later time points.

**Figure 2.**
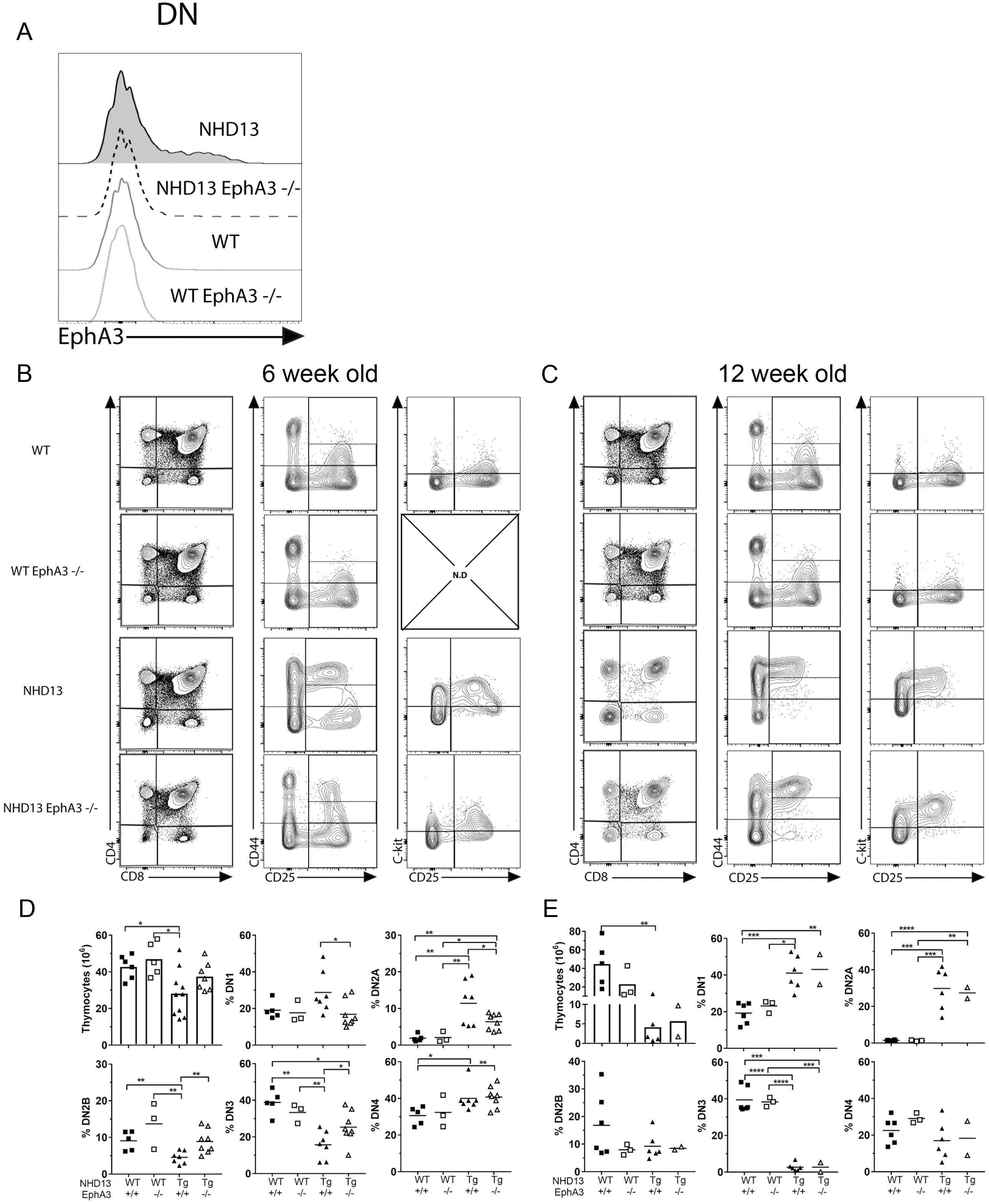
The NHD13 thymocytes differentiation block is transiently rescued by loss of EphA3. **A)** Confirmation of EphA3 knockout in DN (CD4^−^/CD8^−^) thymocytes by flow cytometry. **B)** Representative cell profiles using CD4/CD8, CD44/CD25 and C-Kit/CD25 for each indicated genotype in 6 week old and **C)** 12 week old mice. **D)** Quantitation of thymic cellularity and proportions of DN1, DN2A, DN2B, DN3 and DN4 in 6 week old and **E)** 12 week old mice. Data represent the mean and points represent individual mice 3-8 per group. *P-values* were calculated using Student’s T-test (*<0.05, **<0.005, ***<0.0005). N.D: not done.

### EphA3 is required for long-term but not short-term engraftment

The NHD13 thymocytes abnormal ability to self-renew is functionally defined by their capacity to engraft upon transplantation, and to retain this ability indefinitely upon serial transplantation. This capacity requires the transcription factor Lyl1, yet NHD13-Lyl1^−/−^ mice still develop T-ALL [21]. This self-renewal capacity is therefore not necessary for transformation to occur, and is not the only mechanism by which NHD13 thymus result in the development of T-ALL.

To assess the engraftment ability of NHD13 thymocytes in the absence of EphA3, we performed serial transplantation of NHD13-EphA3^−/−^ thymocytes into sub-lethally irradiated wild type mice, and evaluated donor chimerism 4 weeks later (Fig. 3A). We found that NHD13-EphA3^−/−^ thymocytes are able to engraft similarly to NHD13-EphA3^+/+^ thymocytes (Fig. 3B) with a donor contribution in the DN compartment of 70% and 65% in the first and second transplant, respectively (Fig. 3C). Strikingly, the NHD13-EphA3^−/−^ thymocytes fail to engraft upon the third serial transplant, whereas the NHD13-EphA3^+/+^ thymocytes engraft indefinitely (Fig. 3B and C). In parallel, the cellularity of the thymus (Fig. 3D) and the proportions of the DN1-4 populations (Fig. 3E) were rescued by the third transplant in the NHD13-EphA3^−/−^ recipients and were similar to the WT-EphA3^+/+^ and WT-EphA3^−/−^ control recipients. These experiments were performed on three occasions from independent donors and similar results were seen in each replicate. Therefore, EphA3 is dispensable for short-term engraftment, but essential for long-term engraftment.

**Figure 3.**
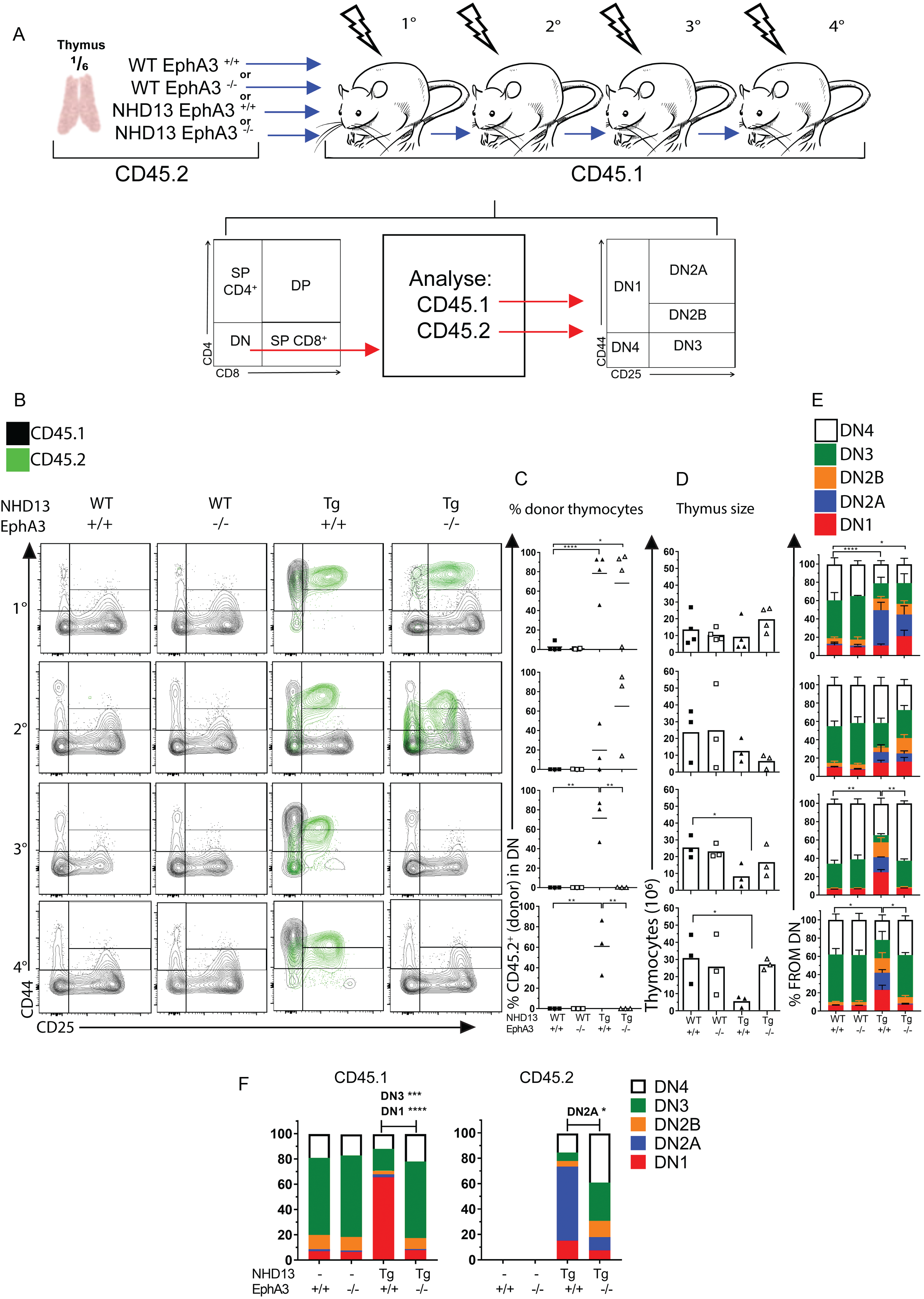
EphA3 mediates the long-term self-renewal capacity of NHD13 thymocytes upon serial transplantation. **A)** Schematic representation of experimental design using 6 week old donor CD45.2 thymi from WT-EphA3 ^+/+^, WT-EphA3 ^−/−^, NHD13-EphA3 ^+/+^ or NHD13-EphA3 ^−/−^ mice. CD45.2 thymocytes equivalent to a ⅙ of a thymus were i.v. injected into sub-lethally irradiated (6.5 Gy) C57BL/6 CD45.1 recipient mice for primary or further serial transplantations. At 4 weeks post-transplantation, engraftment and contribution of donor CD45.2 cells were analysed by flow cytometry. **B)** Representative cell profiles of serial transplants (e.g 1° primary, 2° secondary, 3° tertiary and 4° quaternary) for each indicated genotype show DN1-4 populations determined by CD44/CD25. **Black** plots represent recipient CD45.1 cells and **green** plots represent engrafted donor CD45.2 cells. **C)** Contribution of CD45.2 donor thymocytes of each serial transplant. **D)** Thymus cellularity of each serial transplant. **E)** Quantitation of DN1 to DN4 proportions of each serial transplant. Three biological replicates were performed and the data represents one biological replicate. **F)** Quantitation of DN1 to DN4 split by CD45.1 or CD45.2. C and D) Data represent the mean, points represent individual mice 3 per group and *P-values* were calculated using Student’s T-test (*<0.05, **<0.005, ***<0.0005). E and F) Bars represent the mean ± S.E.M, *P-values* were calculated using a 2way ANOVA and the significance refers to the DN2A populations.

Furthermore, in the primary transplants that received NHD13-EphA3^−/−^ thymocytes, the recipient WT thymocytes (CD45.1) exhibit a strikingly different DN cell profile compared to the thymuses transplanted with NHD13-EphA3^+/+^ thymocytes (Fig. 3B). In the NHD13-EphA3^+/+^ primary recipient, it appears that the WT recipient thymocytes are prevented from differentiating beyond the DN1 stage; there is a dramatic accumulation of the recipient thymocytes in the NHD13-EphA3^+/+^ primary recipient compared to either the WT or EphA3^−/−^ thymuses (Fig. 3B). However, despite the similar level of engraftment of donor cells in the NHD13-EphA3^−/−^ recipient, the WT recipient cells (CD45.1) show a relatively normal DN distribution (Fig. 3F). These results suggest that, upon transplantation of NHD13 thymocytes, EphA3 is required to prevent normal differentiation of the incoming WT progenitor cells.

### NHD13 DN2 thymocytes induce cell cycle arrest on WT cells independently of EphA3 expression

Another characteristic of NHD13-EphA3^+/+^ thymocytes is their increased quiescent state compared to WT cells [21]. This has not yet been assessed in transplant recipients, nor in the context of EphA3. To investigate if EphA3 modulates the cell division of NHD13 thymocytes, we analysed the cell cycling status of DN2 (DN2A and DN2B) thymocytes in the primary transplant recipients.

We found that, following transplant, NHD13-EphA3^+/+^ thymocytes are more quiescent than recipient thymocytes from control WT-EphA3^+/+^ recipient mice (in which no donor cells engraft) (Fig. 4). This is similar to what was previously described for NHD13 thymocytes prior to transplantation [21]. However, there were no differences in the cell cycle status in the NHD13-EphA3^−/−^ cells compared to the NHD13-EphA3^+/+^ cells (Fig. 4), indicating that EphA3 does not affect the cell cycle status of NHD13 thymocytes. Notably, we observed that in both NHD13-EphA3^+/+^ or NHD13-EphA3^−/−^ transplants, the WT-recipient cells (CD45.1) divided at a lower rate compared to the equivalent WT-recipient cells (CD45.1) in the thymuses transplanted with WT-EphA3^+/+^ or WT-EphA3^−/−^ thymocytes (Fig. 4). This observation suggests that, independently of EphA3 expression, NHD13 thymocytes hamper the cell division of proximal WT thymocytes.

**Figure 4.**
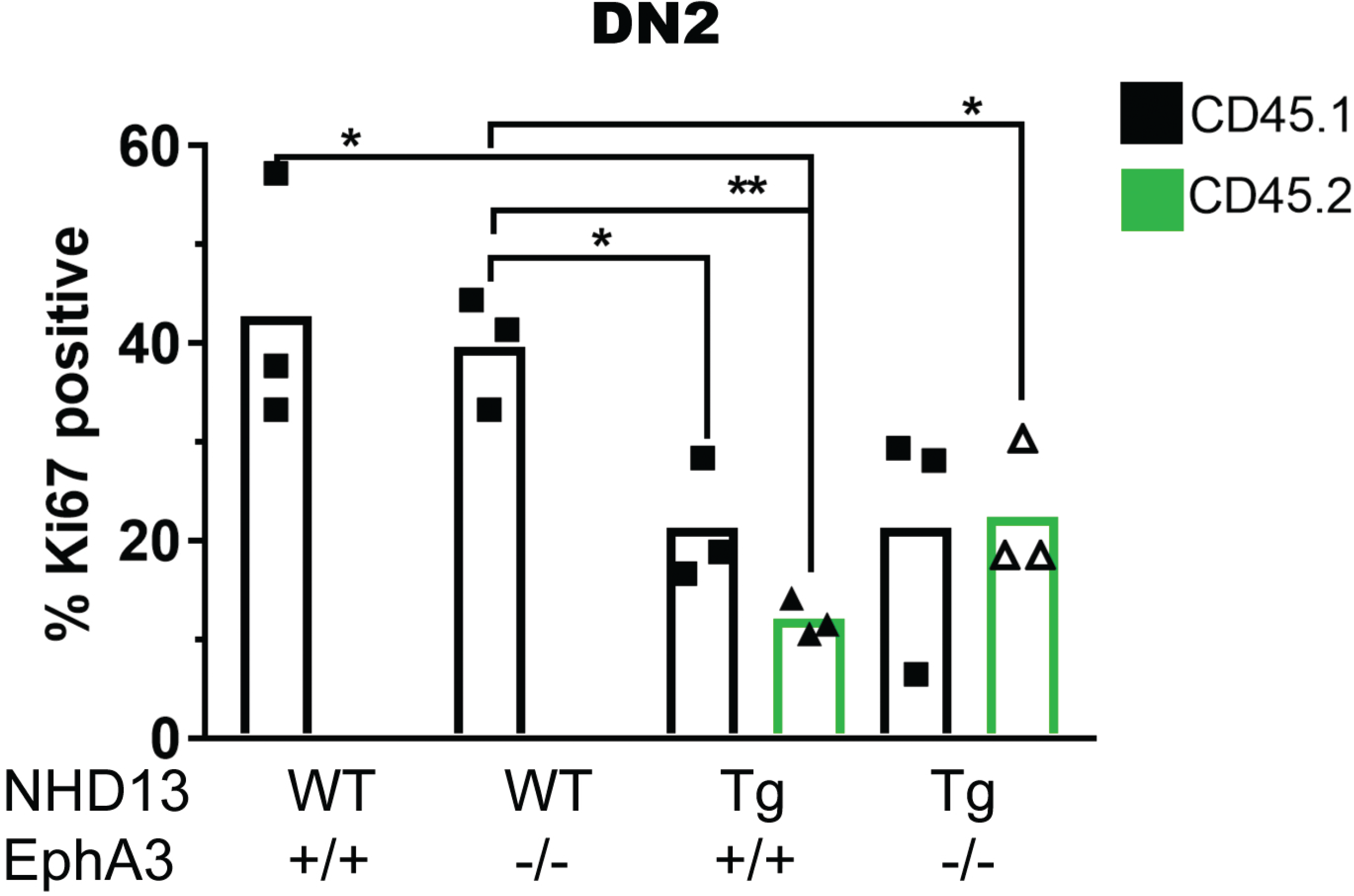
Cell division arrest of WT thymocytes by NHD13 thymocytes. Cell divison analysis of DN2 (DN2A and DN2B) thymocytes in the primary transplant recipient. DN2 (CD44^+^, CD25^+^) recipient (CD45.1) or donor (CD45.2) thymocytes were analysed for cell division/ quiescent status using Ki67 and DNA (Hoechst) stains. Bars represent the mean, points represent individual mice, 3 per group, and *P-values* were calculated using Student’s T-test (*<0.05, **<0.005, ***<0.0005).

### Sustained abnormal expression of self-renewal genes in the absence of EphA3

The stem-like transcriptional signature downstream of the NUP98-HoxD13 fusion protein has LMO2-like, Lyl1 dependent, and Lyl1 independent components [21]. To determine any impact of EphA3 deletion on this self-renewal signature, we compared the expression of several candidate self-renewal regulatory genes in NHD13-EphA3^+/+^ and NHD13-EphA3^−/−^ thymocytes by qRT-PCR. We previously showed overexpression of *HoxA* genes such as *HoxA5, HoxA7, HoxA9* and *HoxA10* in NHD13 DN thymocytes [19]. We found no difference in the profile of overexpression of these genes in whole NHD13 thymus tissue in the presence or absence of EphA3 (Fig.5). Similarly, oncogenes such as *Lmo2, Lyl1, Hhex* and *Klf2*, which are all deregulated in NHD13 DN2 thymocytes, showed no significant changes in expression when comparing NHD13-EphA3^+/+^ and NHD13-EphA3^−/−^ DN2 thymocytes (Fig. 5). These results suggest that EphA3 exerts influence via an independent mechanism to those previously described in T-ALL mouse models.

**Figure 5.**
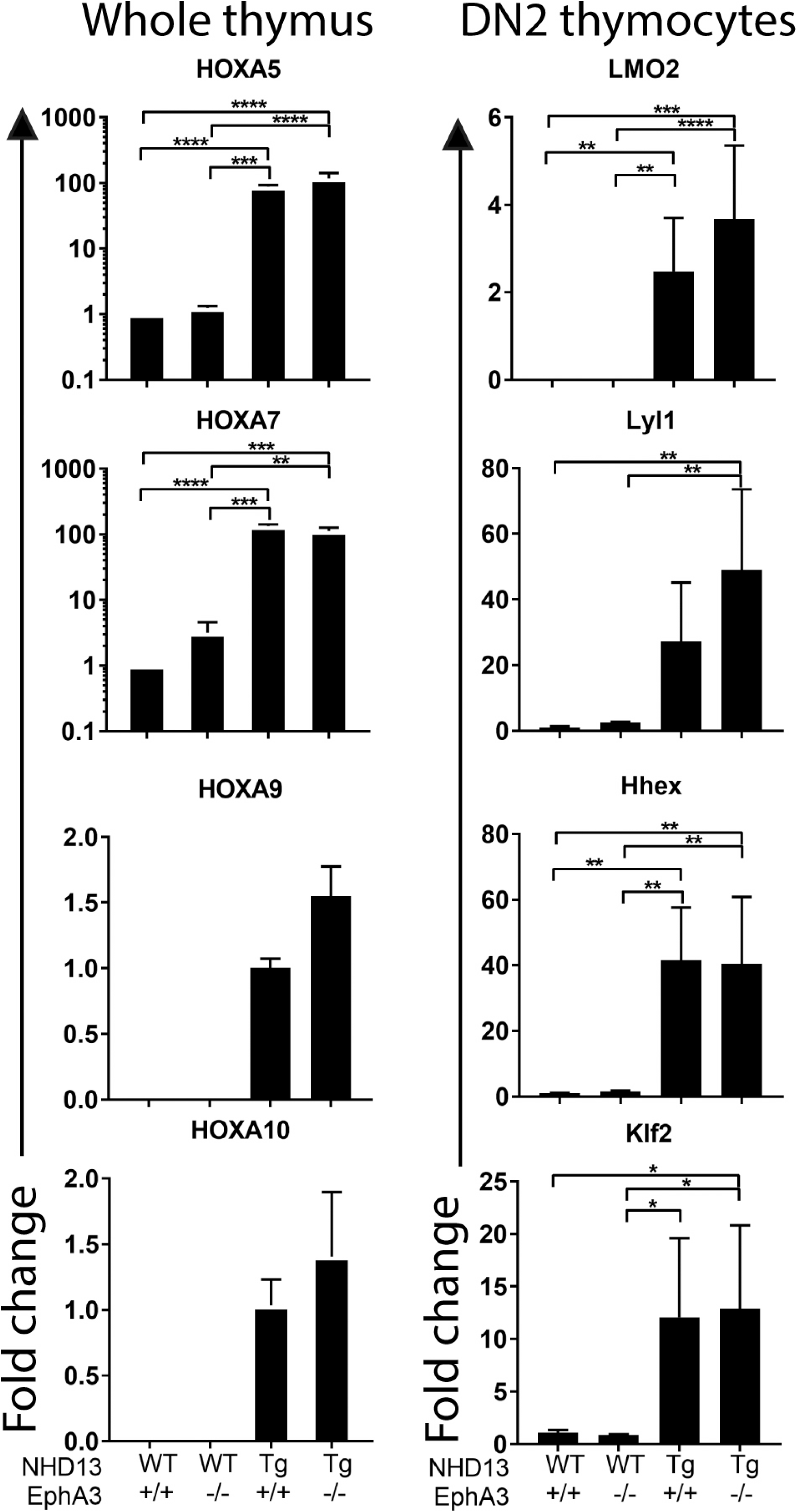
Self-renewal genes. RT-PCR were performed using 6 week old whole thymus (left column) or FACS sorted DN2 (DN2A and DN2B, CD44^+^, CD25^+^ right column) thymocytes samples for each indicated genotype. Average fold change expression of *HoxA5, HoxA7, HoxA9, HoxA10*, *LMO2, Lyl1, Hhex* and *Klf2* genes. Bars represent the mean ± S.E.M of 3 mice per group and *P-values* were calculated using Student’s T-test (*<0.05, **<0.005, ***<0.0005).

### EphA3 is not essential for T-ALL in NHD13 mice

Our studies show that the aberrant expression of EphA3 in the NHD13 thymocytes is responsible for an apparent block on incoming thymocytes, and therefore may contribute to the development of T-ALL in this model by reducing the effectiveness of normal cell competition. To directly determine the impact of the absence of EphA3 on T-ALL occurrence in this model, we compared the T-ALL-free survival of NHD13-EphA3^+/+^ and NHD13-EphA3^−/−^ mice. Both wild type (n = 13) and EphA3^−/−^ (n = 17) cohorts survived without development of any T-ALL. T-ALL occurred in the NHD13-EphA3^−/−^ mice at a lower frequency than in NHD13-EphA3^+/+^ mice (9/30 (30%) NHD13-EphA3^+/+^ mice developed T-ALL, as opposed to 4/25 (16%) NHD13-EphA3^−/−^ mice). The survival curves were significantly different by Mantel-Cox test (p = 0.0418). The curves overlap early in the disease course and then deviate late, perhaps suggesting that EphA3 is especially important in late-onset T-ALL in this model.

We compared the cell profiles of several T-ALLs arising from NHD13-EphA3^+/+^ or NHD13-EphA3^−/−^ mice (Fig. 6B). No differences were noted in the CD4/CD8 profiles between groups, as they vary considerably from sample to sample regardless of the presence of EphA3. Notably, the DN (CD44/CD25) profiles from the NHD13-EphA3^+/+^ show an accumulation of thymocytes in the DN1 (4 of 5) and DN2A (3 of 5) (Fig. 6B), whereas in the absence of EphA3 the DN (CD44/CD25) profiles show thymocytes in other differentiation stages other DN1, including DN3 and DN4. Notably, only one of the five EphA3^+/+^ T-ALLs demonstrated appreciable EphA3 expression. Together, this study demonstrated that deletion of EphA3 reduced but did not eliminate the incidence of T-ALL in the NHD13 model.

**Figure 6.**
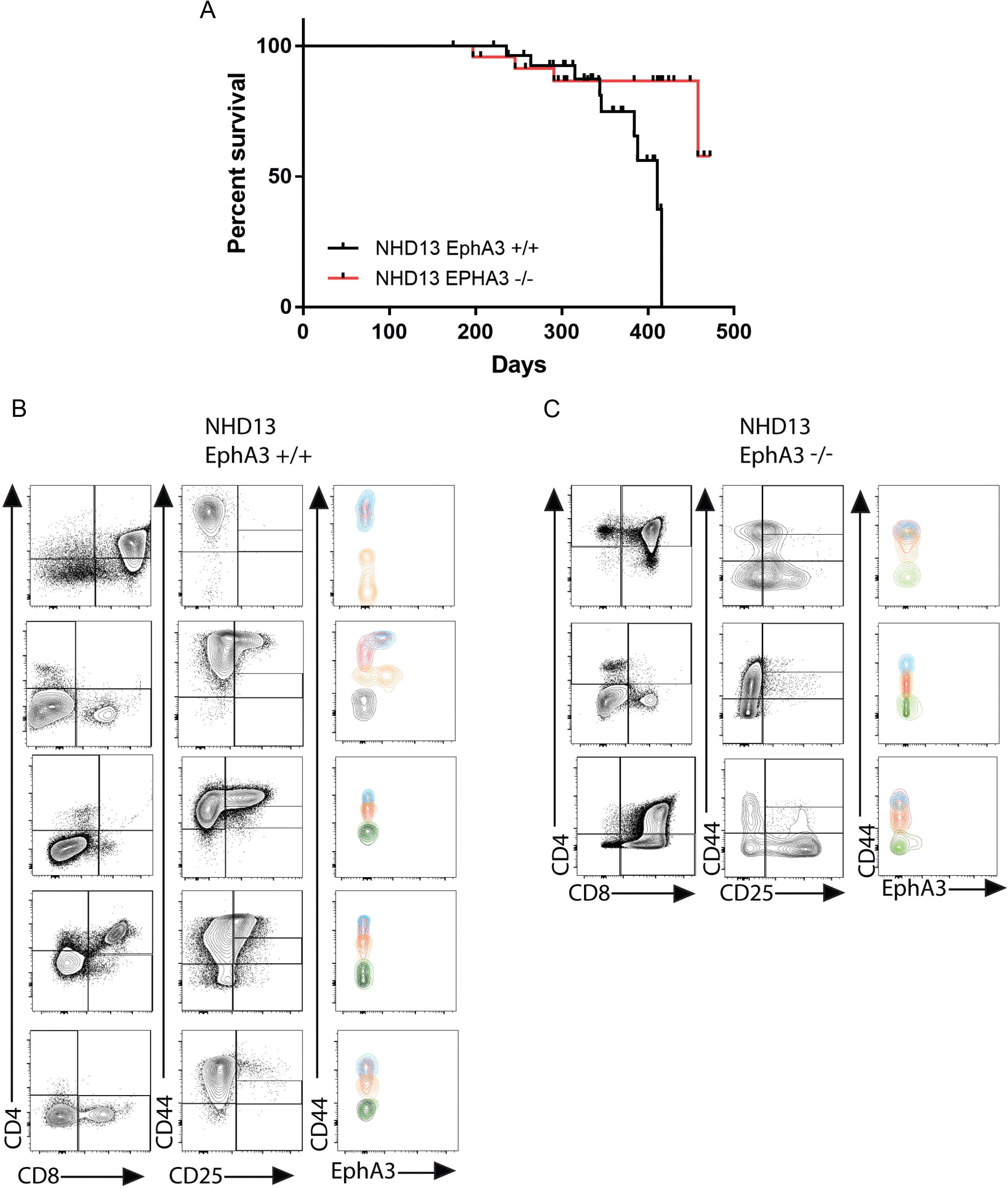
Survival cohort and T-ALL profiles. **A)** Kaplan-Meier plot of T-ALL-free survival for each indicated genotype. **B)** Representative cell profiles and EphA3 detection in T-ALL leukaemias of each indicated genotype using CD4/CD8, CD44/CD25 and CD44/EphA3. The colour of the contour plots matches the colour of the gated DN1-4 populations from figure 1.B.

## Discussion

Pre-leukemic stem cells have previously been characterised on the basis of their ability to self-renew inappropriately, as defined by an engraftment assay. This has been considered a cell-intrinsic ability, granted by the inappropriate expression of stem-like transcription factors. However, the removal of this transplantation ability from NHD13 thymocytes, by deletion of Lyl1, does not impact on T-ALL incidence. Therefore there must be another mechanism responsible.

In the present study, we characterise the role of EphA3 in T-ALL in the NHD13 mouse model. EphA3 is typically considered to be undetectable in adult tissues (both mouse and human), and has been described extensively as overexpressed in a wide variety of haematological malignancies [33, 44, 55]. In myeloid leukemias and glioblastomas, EphA3 has been identified as a regulator of stem cell abilities [46, 47].

We show that EphA3 is not required for the engraftment capacity of pre-leukemic NHD13 thymocytes. These thymocytes have been recently described as a transplantable population of expanded DN2 cells in the NHD13 thymus [21]. Deletion of EphA3 from the NHD13 mice does not prevent the abnormal accumulation or transplantation ability of these cells. However, upon transplantation, these cells are unable to block the normal progression of recipient WT progenitor cells through the normal developmental pathway. This is in contrast to the EphA3^+/+^ NHD13 thymocytes, which block progression of incoming WT progenitors past the DN1 stage.

Previous studies have demonstrated that continual import of progenitors from the bone marrow into the thymus is essential to maintain cell competition in the thymus, a key process to sustain normal differentiation and homeostasis of thymocytes. The absence of normal cell competition results in the abnormal self-renewal of WT thymocytes and is leukemogenic [24–27]. In the models used in the studies which demonstrated this, the mice are deficient in incoming progenitors, so the incumbent thymocytes are not outcompeted and can instead remain in the thymus. This long occupancy presumably permits them time to accumulate mutations and become leukemogenic. In our transplantation model, following the four week engraftment period, recipient thymuses contain two populations of cells: NHD13 cells that were transplanted and engrafted in the thymus, and wild type “progenitor” cells which are newly arrived from the bone marrow. At 4 weeks post transplantation, incoming WT cells accumulate in the DN1 stage as they are prevented from progressing through their normal differentiation pathway by the presence of NHD13-EphA3^+/+^ DN2 cells. This results in an inefficient cell competition in DN3, with no “young” progenitors coming in to outcompete the incumbent “old” progenitors. When EphA3 is deleted, however, transplanted NHD13-EphA3^−/−^ cells still accumulate in the DN2 stage but the incoming WT cells complete their differentiation process normally, restoring cell competition in DN3. Consistent with this, the cellularity of the thymus is restored, and over serial transplants, while the NHD13-EphA3^+/+^ cells engraft repeatedly, the NHD13-EphA3^−/−^ cells fail to engraft in the third transplantation. This phenomenon may explain the differences seen between non-transplanted NHD13-EphA3^+/+^ and NHD13-EphA3^−/−^ thymuses at six weeks of age but which were absent at twelve weeks of age (Figure 2); the progression of incoming progenitors beyond DN1 in the absence of EphA3 may be more normal than in the presence of EphA3, but because these incoming progenitors are still expressing NHD13, they mostly accumulate in DN2 and predominantly do not progress normally. This is different to the transplant setting, where the incoming progenitors are wild type recipient cells, and will progress normally if they are able.

Based on the extracellular nature of EphA3 and the fact that DN2 NHD13-EphA3^+/+^ thymocytes are known to be responsible for the impaired thymocyte turnover [21], it seems likely that EphA3 may be mediating a cell-cell interaction that stops the incoming WT thymocytes in the DN1 stage, potentially killing them, or blocking them in DN1 and preventing new progenitors from importing into the thymus behind them.

This “blockade” capacity is independent of the ability of NHD13 thymocytes to accumulate in the DN2 stage, as this accumulation occurs in the absence of EphA3. This accumulation is likely driven by the maintained self-renewal gene expression signature of HoxA and Lyl genes. Another potentially concurrent possibility is that the accumulation of DN2 thymocytes is dependent on the abnormal expression of c-kit. Stem cell factor (SCF) is the ligand of c-kit and is a chemokine released by the thymic microenvironment in the DN1 and DN2 niches [56, 57]. Notably, c-kit signalling is required for the transplantation ability of Lmo2 transgenic thymocytes [58]. Here we report that the DN2 NHD13 thymocytes overexpress c-kit and accumulate independently of EphA3 expression. Thus, we propose that in this model the DN2 accumulation of NHD13 thymocytes is an ability distinct from the prevention of normal thymocyte cell competition, only the latter of which is mediated by EphA3.

In the NHD13 mouse model, the ablation of the cell competition inhibition mechanism by deletion of EphA3 is insufficient to prevent the development of T-ALL (although it does reduce the incidence). Cell competition is restored by deletion of EphA3, but the intrinsic self-renewal capacity of the NHD13 thymocytes driven by LMO2/Lyl1 [21] remains intact. Although Lyl1 is required for the transplantability and establishment of a stem cell-like gene expression program in NHD13 thymocytes, deletion of Lyl1 is also not sufficient to prevent T-ALL [21]. Therefore, we suggest that development of T-ALL in this model is a complex interplay of cell intrinsic and cell extrinsic factors. It is clear that NHD13 thymocytes employ at least two different mechanisms of leukemogenesis: 1) the expression of the oncogene Lyl1, and 2) the inhibition of cell competition mediated by EphA3.

Our findings also report insights into new cellular mechanisms by which NHD13 pre-leukemic stem cells are able to manipulate WT normal cells. We found that both NHD13 and wild type cells in recipient thymuses divide at a slower rate. This suggests that the presence of the NHD13 cells can inhibit the normal cell division of the wild type cells. This effect was independent of EphA3, indicating that there is more than one means of communication between NHD13 and wild type thymocytes in our transplant model.

In the present study, deletion of EphA3 resulted in the restoration of normal cell competition in NHD13 thymuses, and reduced the incidence of T-ALL as a result. This supports earlier work [25] demonstrating that cell competition is a tumour suppressor in the thymus, in a model which was not specifically engineered to disrupt incoming thymocyte progenitors. We have, for the first time, identified a cell competition mechanism in a non-engineered model, and further identified a molecular mediator responsible for the establishment of this mechanism. Given the perceived role of EphA3 in self-renewal in other settings, it is interesting to ponder whether these phenomena are also due to a contribution of cell competition to self-renewal.

## Acknowledgements

We thank the UQ Biological Resources facility at the Translational Research Institute, Brisbane, Australia, and the Flow Cytometry facility at the Translational Research Institute, Brisbane, Australia, for housing our experiments. We also thank J. Daniel Bautista for providing the mouse illustrations. This work was funded by NHMRC (Project Grant APP1099381) and Advance Queensland, Women’s Academic Fund.

## Author Contributions

A. Pliego Zamora designed research, performed most experiments and analysis, and wrote the manuscript. H. Ranasinghe, J. Lisle, S. Huang and R. Wadlow performed additional experiments. A. Scott provided the IIIA4 antibody. A. Boyd designed research and provided expertise regarding EphA3 function. C. Slape designed research, performed additional experiments, analysed data and wrote the manuscript. All authors approved the final draft.

---

**Supplementary figure 1.**
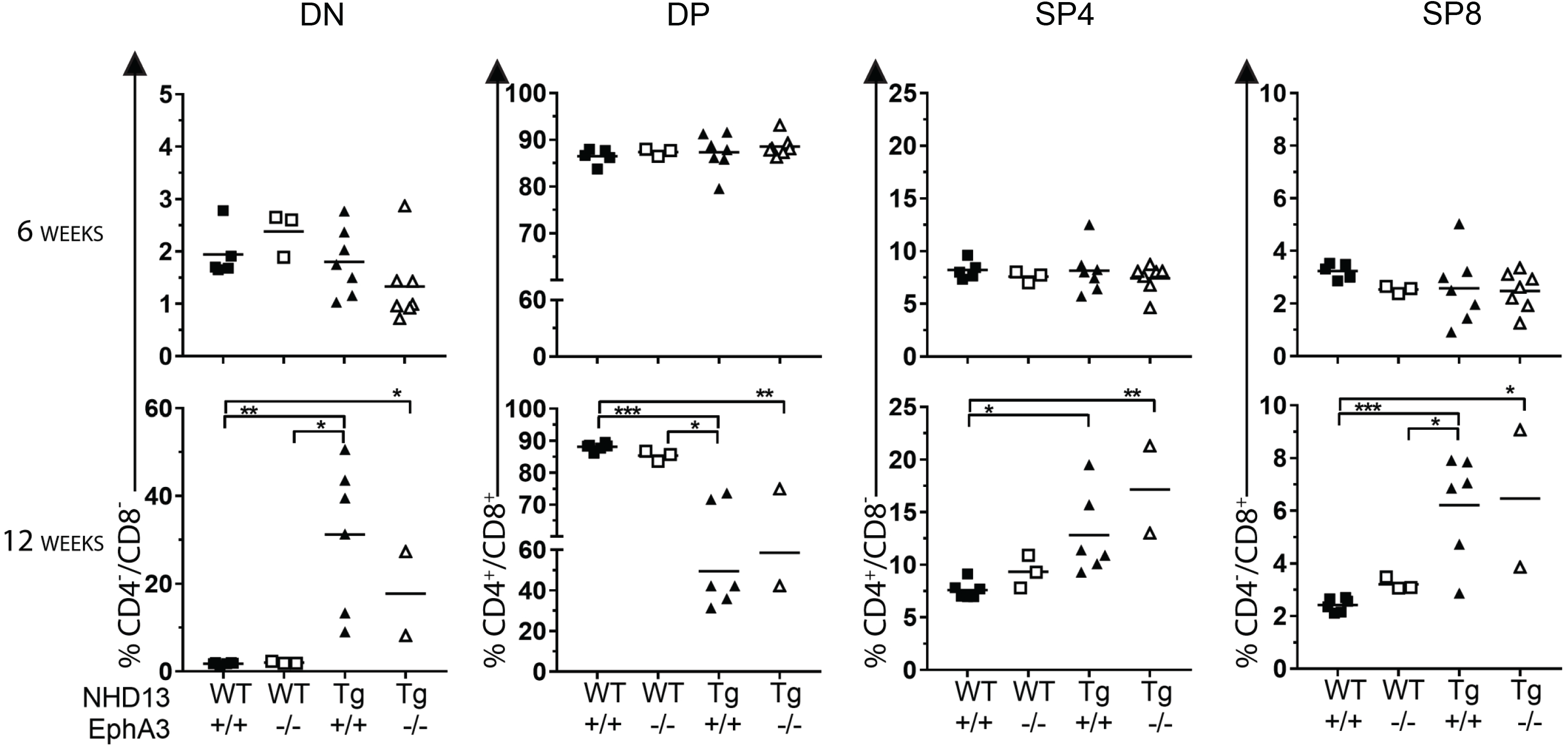
Quantitation of Double Negative (DN), Double Positive (DP), Single Positive CD4+ (SP4) and Single Positive CD8+ (SP8) populations. Quantitation of DN (CD4^−^/CD8^−^), DP (CD4^+^/CD8^+^), SP4 (CD4^+^/CD8^−^) and SP8 (CD4^−^/CD8^+^) in 6 and 12 week old mice of each indicated gentoype. Data represent the mean and points represent individual mice 3-8 per group. *P-values* were calculated using Student’s T-test (*<0.05, **<0.005, ***<0.0005).

